# Capped nascent RNA sequencing reveals novel therapy-responsive enhancers in prostate cancer

**DOI:** 10.1101/2022.04.08.487666

**Authors:** Kellie A. Cotter, Sagar R. Shah, Mauricio I. Paramo, Shaoke Lou, Li Yao, Philip D. Rubin, You Chen, Mark Gerstein, Mark A. Rubin, Haiyuan Yu

## Abstract

Mounting evidence suggests that enhancer RNA (eRNA) transcription start sites (TSSs) provide higher sensitivity and specificity for enhancer identification than histone modifications and chromatin accessibility. The extent to which changes in eRNA transcription correspond to changes in enhancer activity, however, remains unclear. Here, we used precision run-on and capped RNA sequencing (PRO-cap) to assess changes in enhancer activity in response to treatment with the androgen receptor signaling inhibitor, enzalutamide (ENZ). We identified 6,189 high-confidence candidate enhancers in the human prostate cancer cell line, LNCaP; 853 of which demonstrated significant changes in activity in response to drug treatment. Notably, we found that 67% and 54% of drug-responsive enhancers did not show similar changes in activity in previous studies that utilized ChIP-seq and ATAC-seq, respectively. Strikingly, 79% of regions with increased eRNA transcription showed no other biochemical alterations, implying that PRO-cap can capture a set of precise changes in enhancer activity that classical approaches lack the sensitivity to detect. We performed *in vivo* functional validations of candidate enhancers and found that CRISPRi targeting of PRO-cap-specific drug-responsive enhancers impaired ENZ regulation of downstream target genes, suggesting that changes in eRNA TSSs mark true biological changes in enhancer activity with high sensitivity. Our study highlights the utility of using PRO-cap as a complementary approach to canonical biochemical methods for detecting precise changes in enhancer activity and, in particular, for better understanding disease progression and responses to treatment.

## Main

First-line treatment for advanced metastatic prostate cancer (PCa) generally involves androgen deprivation therapy (ADT) to reduce the activity of androgen receptor (AR)(Heinlein and Chang, 2004). Of particular clinical concern is metastases of castration-resistant PCa (CRPC), where the disease has developed resistance to both first-line and second-generation AR signaling inhibitors (ARSi, e.g., enzalutamide)(Scher and Sawyers, 2005). Thus, determining both the mechanisms behind ADT resistance and the distinct signaling pathways activated in CRPC are essential to improving existing therapies and finding new potential drug targets.

Given the enrichment of genomic alterations in metastatic PCa(Armenia et al., 2018), there are increased efforts to understand the complexity of the PCa genome. Recent studies using whole genome sequencing have revealed several examples of alterations in gene regulatory regions. For example, upwards of 80% of samples were found to have a duplication of a region upstream of *AR*, which was then shown to be a previously unidentified enhancer(Quigley et al., 2018; Takeda et al., 2018; Viswanathan et al., 2018). This enhancer duplication was shown to increase the expression of *AR* and in doing so decrease sensitivity to ARSi. Therefore, it is important to precisely identify and characterize enhancer dynamics in response to therapeutic intervention in PCa. However, despite the exhaustive amount of sequencing information captured by WGS, our interpretation and understanding of the data is limited due to the incomplete annotation of the non-coding genome, including transcriptional regulatory elements such as enhancers.

In general, enhancer regions are currently defined by biochemical features such as chromatin accessibility (DNase I hypersensitivity or transposase accessibility) and histone modification marks (H3K27ac and H3K4me1) along with transcription factor binding profiles as determined by ChIP-seq(Gasperini et al., 2020). Large-scale reporter assays (e.g., STARR-seq or MPRA) have also been used to evaluate the enhancer potential of candidate DNA regions(Arnold et al., 2013; Kheradpour et al., 2013). However, these reporter assays have consistently shown that less than 50% of regions with these biochemical annotations exert enhancer activity(Kwasnieski et al., 2014; Vanhille et al., 2015). Likewise, similar techniques have been used to identify thousands of potential enhancer regions in PCa by combining ChIP-seq and whole genome STARR-seq(Kron et al., 2017; Liu et al., 2017; Sharma et al., 2013; Stelloo et al., 2018). In addition, chromosome capture has been used to elucidate chromatin interactions genome-wide and those loci specifically associated with AR and RNA Polymerase II using ChIA-PET(Ramanand et al., 2020; Rhie et al., 2019; Taberlay et al., 2016; Zhang et al., 2019). Interestingly, a recent study demonstrated that the majority of AR binding sites are not active enhancers(Huang et al., 2021). Moreover, despite increased recruitment of AR to these regions, dihydrotestosterone (DHT) stimulation did not increase enhancer activity in over half of the AR-bound active enhancers. Perhaps most intriguing, however, CRISPR interference of some “inactive” enhancers altered the expression of the enhancer-regulated gene, at times at a similar level of interference to nearby “active” enhancers. Overall, this highlights the limitations of the datasets produced thus far and the need for alternative methods for enhancer identification given that epigenomic-mark-based approaches identify enhancers with a high false-positive rate, while reporter assays cannot fully reproduce much of the biological complexity of large enhancers regions in their native genomic context, in particular with regards to multiple enhancers acting on a single gene.

More recently, widespread RNA polymerase II-mediated bidirectional transcription has been observed in enhancer regions, which produce biochemically unstable transcripts known as enhancer RNAs (eRNAs)(Core et al., 2014; Kim et al., 2010; Tippens et al., 2018). Although the functional significance of eRNAs remains unclear, evidence suggests that enhancer transcription corresponds with activation(Chen et al., 2018; Chen and Liang, 2020), with close to 50% of short capped nascent RNAs that map to previously unannotated TSSs overlapping with episomal reporter-validated enhancers(Henriques et al., 2018). Moreover, it has been found that ∼95% of putative active enhancers found within accessible chromatin drive local transcription and do so using factors and mechanisms overwhelmingly similar to those of promoters(Core et al., 2014). Recently, we performed systematic interrogation of enhancer elements and showed that active enhancer units are precisely marked by divergent eRNA TSSs genome-wide. Moreover, although eRNA transcription is closely correlated with histone marks, we saw that these epigenomic marks offer lower resolution and specificity for enhancer activation than transcription initiation(Tippens et al., 2020). Thus, these data support a model whereby transcription is required for distal enhancer function, challenging classical methods that rely on chromatin accessibility and histone modifications to identify active enhancers.

On average, enhancers transcribe at 5% the level of promoters(Core et al., 2014; Henriques et al., 2018). Thus, due to their low abundance and instability, eRNA detection requires highly sensitive alternative methods to standard RNA sequencing approaches. Recently, we performed systematic comparisons of genome-wide RNA sequencing assays suitable for the identification of active enhancers and found that the nuclear run-on followed by cap-selection assays (namely, Global/Precision Run-On and capped RNA sequencing GRO/PRO-cap) provide the highest sensitivity and specificity for eRNA detection and active enhancer identification across the whole genome(Yao et al., 2022). Importantly, PRO-cap libraries undergo a series of cap state selection reactions to modify the 5′ ends of transcripts and allow for the accurate identification of transcription initiation sites (Figure 1A). As a result, PRO-cap is highly sensitive for capturing eRNA transcription and therefore, a powerful tool for enhancer identification.

**Figure 1.**
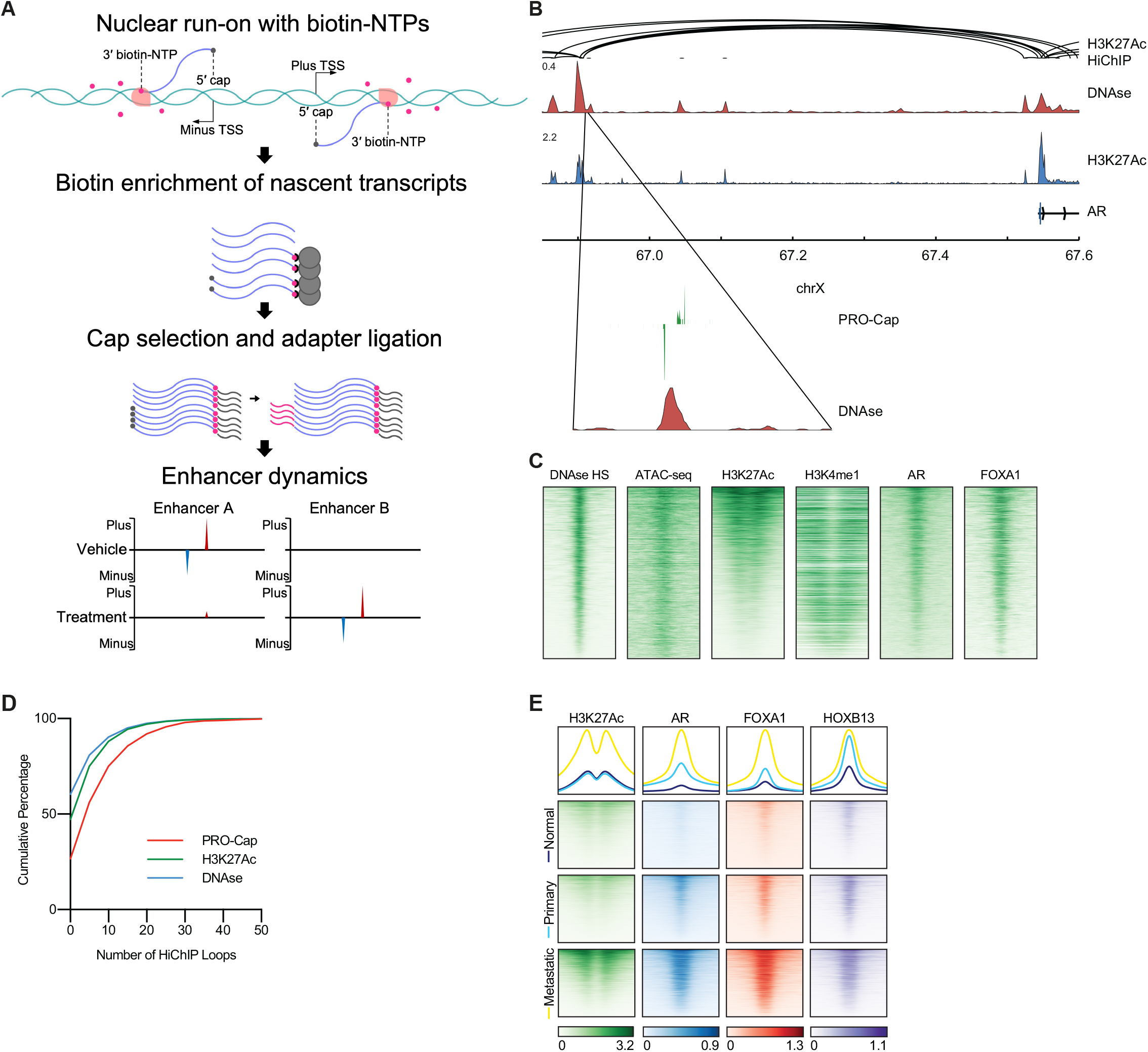
PRO-cap sequencing of eRNAs to map active enhancers in prostate cancer. **A)** Overview of experimental pipeline implemented in this study. **B)** Enhancer candidates demonstrate divergent transcription of eRNAs. Shown is the upstream enhancer for androgen receptor (AR) along with published H3K27Ac HiChIP loops(Giambartolomei et al., 2021) and read tracks for DNase (GSM816637) and H3K27Ac (GSE85558) ChIP-seq(Meuleman et al., 2020; Shukla et al., 2017) of LNCaP cells. **C)** Putative enhancers identified by PRO-cap (n = 6,189) are bound by canonical biochemical marks and known important prostate transcription factors in LNCaP cells. Heatmaps show read density from publicly available ChIP-seq data(Kim et al., 2018; McNair et al., 2018; Meuleman et al., 2020; Shukla et al., 2017; Taberlay et al., 2016) (DNase: GSM816637, AR/FOXA1: GSE85558, ATAC-seq: GSE105116, H3K4me1: GSE73783, and H3K27Ac: GSE107780) 1kb up- and downstream of the center of the enhancer. **D)** H3K27Ac loop anchors(Giambartolomei et al., 2021) are significantly more likely (*P* < 0.0001, Kolmogorov-Smirnov test) to overlap with PRO-cap peaks versus H3K27Ac or DNase I peaks. **E)** Putative enhancers identified by PRO-cap are also bound by biochemical marks and prostate transcription factors in patient samples. Heatmaps show average read density from publicly available ChIP-seq(Pomerantz et al., 2020) data (GSE130408) from normal prostate tissue, primary PCa, and metastatic PCa (n = 8 samples for each).

Here, we utilized changes in eRNA expression to assess changes in enhancer activity in response to short-term treatment with the ARSi enzalutamide (ENZ). We applied PRO-cap to enrich and sequence only nascent RNAs associated with engaged RNA polymerase and to identify divergent transcription start sites marking active enhancers at base pair-resolution(Mahat et al., 2016; Tippens et al., 2020). We identified over 6,000 candidate enhancers in LNCaP cells; 853 of which demonstrated significant changes in enhancer activity in response to ENZ treatment. Importantly, these results identified a large percentage of therapy-responsive enhancers, which were not previously shown to be responsive in studies utilizing other biochemical marks (H3K27ac-ChIP-seq/ATAC-seq) or reporter assays (i.e., STARR-seq). Our study highlights the utility of using eRNA transcription, in particular PRO-cap, as a generalizable and complementary approach to canonical biochemical methods for detecting precise changes in enhancer activity, specifically for applications in disease prognosis, progression, and treatment.

## Results

### Identification of active enhancers in a prostate cancer cell line model using PRO-cap

To measure enhancer activity in their native genomic and cellular context, we utilized PRO-cap to preferentially sequence nascent transcriptional start sites including those generating eRNAs (Figure 1A). Overall, this analysis in LNCaP cells identified 91,705 statistically significant peaks (n = 2 replicates), 68.4% of which were in gene distal regions. 6,189 of these gene distal regions contained divergent significant peaks on both the plus and minus strands and were extended by 200 bp in both directions and called as high confidence enhancer regions (Data S1). As expected, the majority of these regions contained canonical biochemical marks delineating enhancers including open chromatin (DNase I hypersensitivity, ATAC-seq), H3K27 acetylation, and H3K4 monomethylation, and are bound by prostate-enriched transcription factors (AR and FOXA1) (Figure 1B-C). However, 11% (n = 674) of these regions did not exhibit those classical biochemical features (at least two of the three: DNase I HS, H3K27ac, or H3K4me1), demonstrating that PRO-cap identifies a novel set of enhancer regions that would otherwise be missed (Figure 1C).

Using publicly available H3K27ac HiChIP data, we estimated the number of chromatin connections of each candidate enhancer with other genomic regions(Giambartolomei et al., 2021). We found that the regions delineated by PRO-cap have a significantly greater number of loops (*P* < 0.0001, Kolmogorov-Smirnov test) than H3K27ac or DNase I peaks (Figure 1D). Next, we inquired whether these enhancers identified in LNCaP cells were expressed in patient-derived PCa samples. To that end, we analyzed published ChIP-seq data from clinical specimens of non-neoplastic prostate, primary PCa, and metastatic PCa(Pomerantz et al., 2020). As seen in Figure 1E, these tissues recapitulate H3K27ac and the binding pattern of prostate-enriched transcription factors (AR, FOXA1, and HOXB13) in enhancers seen in LNCaP cells. Furthermore, given that LNCaP cells are derived from an androgen-responsive metastatic lymph node lesion(Horoszewicz et al., 1983), the enhancer expression profiles are indeed strongest in metastatic PCa tissues. Overall, these results demonstrate that, unlike traditional epigenomic-based methods, PRO-cap can uncover previously unknown enhancer loci with hallmarks of PCa-relevant TF-binding patterns observed in primary and metastatic lesions.

### Measurement of changes in enhancer activity in response to androgen deprivation therapy

Given our confidence in detecting enhancers by PRO-cap, we investigated whether this method could be utilized to assess changes in enhancer activity in response to treatment. To do so, we treated LNCaP cells with the ARSi enzalutamide (ENZ) for 4 or 24 hours. Using the same analysis as described above, we identified 6769 and 8513 high confidence enhancers using PRO-cap in the 4- and 24-hour time point conditions, respectively (Data S2-3). Approximately 4,479 of these enhancers were identified in both ENZ timepoints and 2,922 were identified irrespective of treatment (Figure S1A).

In addition to detecting enhancers that are expressed at only a single treatment time point, we were interested in identifying those which were active at more than one time point, but with differential levels of expression. Thus, we compiled the total list of enhancers found in any of the two treatment timepoints and vehicle and assessed their differential expression upon treatment (versus vehicle) on at least one strand. This analysis identified 853 enhancers that were significantly activated or repressed after 24 hours of ENZ treatment (Figure 2A, Figure S1B-D, Data S4-5). Furthermore, 73 of these regions were significantly activated or repressed on both the plus and minus strands.

**Figure 2.**
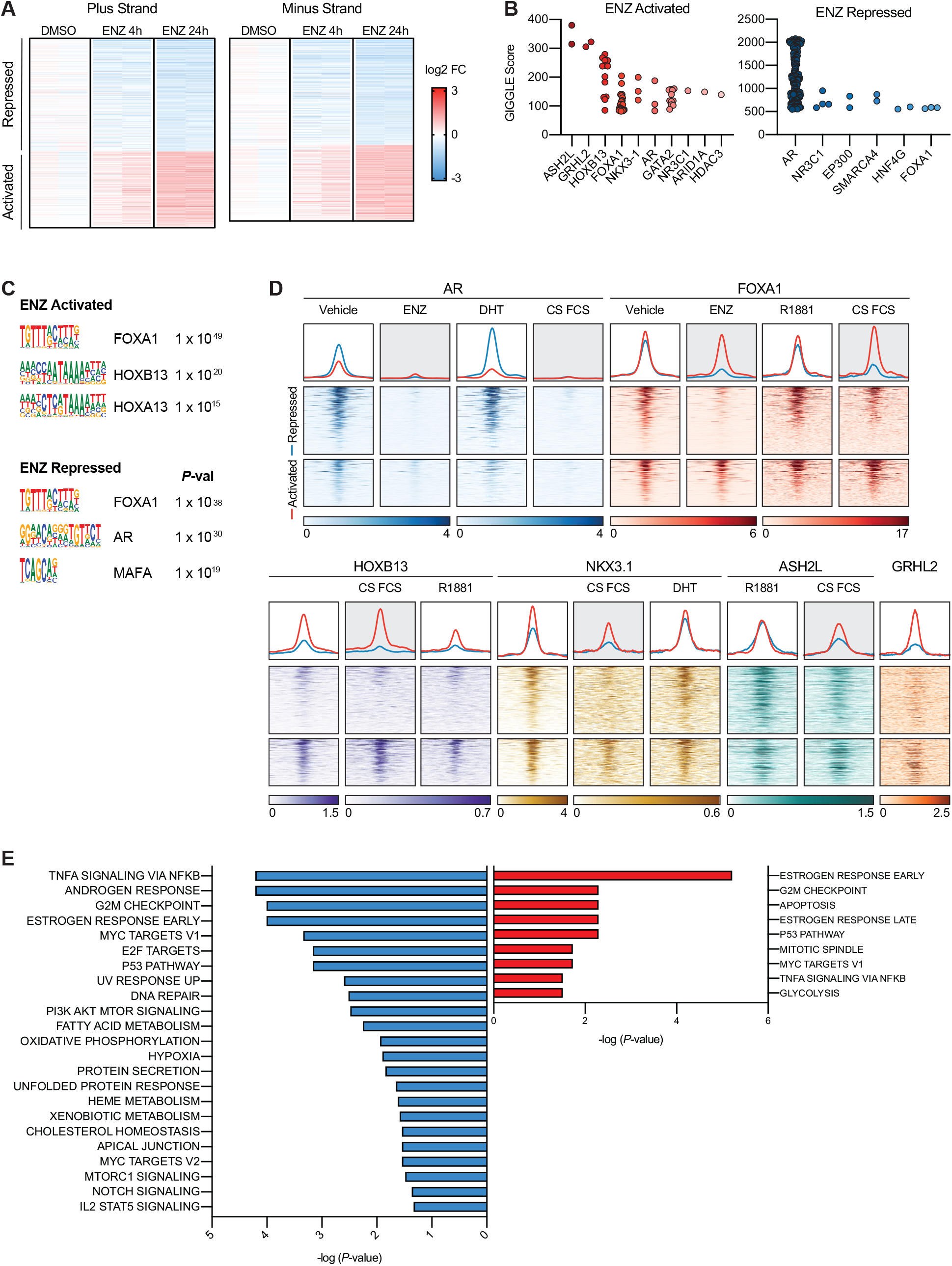
PRO-cap sequencing of eRNAs identifies enhancers affected by enzalutamide. **A)** PRO-cap analysis of LNCaP cells treated with 10 µM enzalutamide (ENZ) for 24 hours identified 853 putative enhancers which demonstrated significant differential activation or repression on the plus or minus strands as determined by edgeR analysis (FDR < 0.05). Data is displayed as a heatmap of the fold-change in mapped reads with 4 or 24 hour ENZ treatments. For each treatment condition two independent biological replicates are shown. **B)** GIGGLE(Layer et al., 2018) analysis demonstrates overlap of the ENZ regulated enhancers with regions identified in published ChIP-seq datasets. Each dot represents a single study, and the GIGGLE score incorporates both enrichment and significance. **C)** Overrepresented TF motifs as determined by Cistrome MDSeqPos(Mei et al., 2017) for the ENZ regulated enhancers. **D)** Heatmaps and summary plots show read density 1kb up- and downstream of the center of the ENZ regulated enhancers. Data is from publicly available ChIP-seq data(Hwang et al., 2019; Palit et al., 2019; Paltoglou et al., 2017; Pomerantz et al., 2015; Rasool et al., 2019; Tan et al., 2012) (GSE137775, GSE125245, GSE94682, GSE70079, GSE40269, GSE28264, and GSE80256) for LNCaP cells treated with ENZ, R1881, DHT or cultured in media containing charcoal stripped (CS) FCS. ASH2L data(Malik et al., 2015) (GSE60841) is from VCaP cells treated with R1881 or cultured in media containing CS FCS. **E)** Gene ontology analysis of the genes predicted to be regulated by these enhancers generated using ShinyGO(Ge et al., 2020).

To predict putative transcription factors (TF) regulating these enhancer regions, we utilized GIGGLE(Layer et al., 2018) analysis (Figure 2B) to probe the CISTROME(Mei et al., 2017) ChIP-seq database containing published datasets with overlapping genomic regions. Of the TFs predicted to bind to the ENZ repressed elements, AR was the top candidate with the highest GIGGLE score. Interestingly, other significant TF predicted to bind the ENZ repressed enhancers included glucocorticoid receptor (*NR3C1*), HNF4G (a TF which drives ARSi resistance(Shukla et al., 2017)), and the co-activator EP300, the SWI/SNF family chromatin remodeler SMARCA4, and the pioneering factor FOXA1. In addition to AR, FOXA1, and NR3C1, other TF predicted to bind the ENZ-activated enhancers included the histone methyltransferase ASHL2, the AR co-regulators GRHL2 and HOXB13, the AR-regulated TF NKX3-1, the pioneering factor GATA2, ARID1A another SWI/SNF family member, and the histone deacetylase HDAC3. Motif analysis further supported these predictions. Motifs for FOXA1, AR, and MAFA were significantly enriched in ENZ-repressed enhancers, while FOXA1, HOXB13, and HOXA13 motifs were significantly enriched in the ENZ-activated enhancers (Figure 2C).

We further mined published ChIP-seq data from LNCaP cells to determine how the binding of the identified enhancer-associated TF is changing in response to ENZ treatment or androgen stimulation with dihydrotestosterone (DHT) or the synthetic androgen, R1881) (Figure 2D). Both ENZ-activated and -repressed enhancers lose AR binding in response to ENZ (AR inhibition) or the absence of hormones (charcoal-stripped, CS FCS), and binding is regained with androgen stimulation. Strikingly, FOXA1 binding at the ENZ repressed enhancers is lost with ENZ or CS FCS treatments and regained with androgen stimulation, while the binding is unaffected at ENZ activated enhancers. Similar patterns were seen with NKX3.1 and ASH2L with dynamic changes in binding seen at the ENZ repressed enhancers with trivial changes at the activated ones. In contrast, both GRHL2 and HOXB13 show minimal binding at the ENZ repressed enhancers as compared to the activated enhancers irrespective of treatment. Similar patterns were seen between the activated and repressed groups with ChIP-seq analysis of metastatic patient samples(Pomerantz et al., 2020) (Figure S1E).

Using a previously described method(Wang et al., 2018), we predicted which genes are regulated by each ENZ-responsive enhancer. We generated predictions for 690 of the 853 regions with an average of 6.3 genes per candidate enhancer (Figure S1F, Data S6-7). GO analysis of these gene lists demonstrated that the majority of genes are related to steroid hormone signaling (in particular androgen response) or the cell cycle (Figure 2E).

Altogether, these findings demonstrate that PRO-cap can detect a large set of precise changes in enhancer activity that other approaches lack the sensitivity to capture.

### Non-coding mutation analysis in PRO-cap-detected ENZ-responsive candidate enhancers

We next queried if PRO-cap-detected enhancer regions could help prioritize somatic genomic variation in non-coding regions. To that end, using 286 PCa whole genomes available through ICGC, we searched for single nucleotide variants (SNVs) in the identified 853 ENZ-regulated enhancers. In total 137 variants were discovered in these regions; 20 of which were recurrent in more than one patient (Figure S2A). Interestingly, eight of these enhancers also had more than one SNV within the enhancer region. An example of this is shown in Figure S2B in which two patients have a recurrent SNV which disrupts an ESRRA motif within the enhancer, and two different patients have a recurrent SNV which creates an HNF4 motif within the enhancer.

Next, we investigated whether the PRO-cap-identified enhancers may harbor potential PCa-associated germline single nucleotide polymorphisms (SNPs) (Conti et al., 2021). Of the 269 known PCa risk variants, none overlap with our 853 ENZ-regulated enhancers. However, 10 SNPs did overlap with our larger list of 6,189 total high confidence PCa enhancers (*P* < 0.001). A representative example of a significantly enriched SNP at a PCa enhancer is shown in Figure S2C. It is noteworthy that none of these SNPs have been previously identified as residing in gene regulatory regions. Thus, these results highlight the power of PRO-cap in identifying and delimiting the non-coding regulatory genome to prioritize enhancer-associated mutations.

### *in vivo* functional validation of PRO-cap-detected ENZ-responsive candidate enhancers

Next, we sought to determine how PRO-cap compares with other methods at measuring enhancer activity changes. We first surveyed published H3K27ac ChIP-seq and ATAC-seq data from LNCaP cells treated with ENZ. Many of the regions with altered enhancer activity as measured by PRO-cap could also be detected by changes in the ChIP- or ATAC-seq data (Figure 4A). Surprisingly, however, 67% and 54% of the regions did not show a similar change in the ChIP- or ATAC-seq data, respectively (highlighted in navy and burgundy). Particularly striking was that 78-79% of the regions with an increase in eRNA transcription with ENZ showed no biochemical alterations.

Given our observations, we sought to validate the enhancer activity of our candidates identified by PRO-cap using an *in vivo* approach via CRISPR interference (CRISPRi) of these enhancers. To that end, we genetically targeted two of the previously tested candidates (#1 and #3) that were repressed by ENZ (Figure 4B). Enhancer candidate #1 is located within in intronic region of *CACNG4*, while enhancer candidate #3 is found within the intron of *KCNMA1*, (Figures S3-4). Both of these genes are known to be repressed by ENZ treatment. We designed 3 sgRNAs each against these regions centered around the TSS of the eRNAs. We then transfected dCas9-KRAB-stably expressing LNCaP cells with these sgRNAs. CRISPRi-mediated repression of the two candidate enhancers reduced the expression of both the eRNA and the predicted target genes (Figure 3C-D).

**Figure 3.**
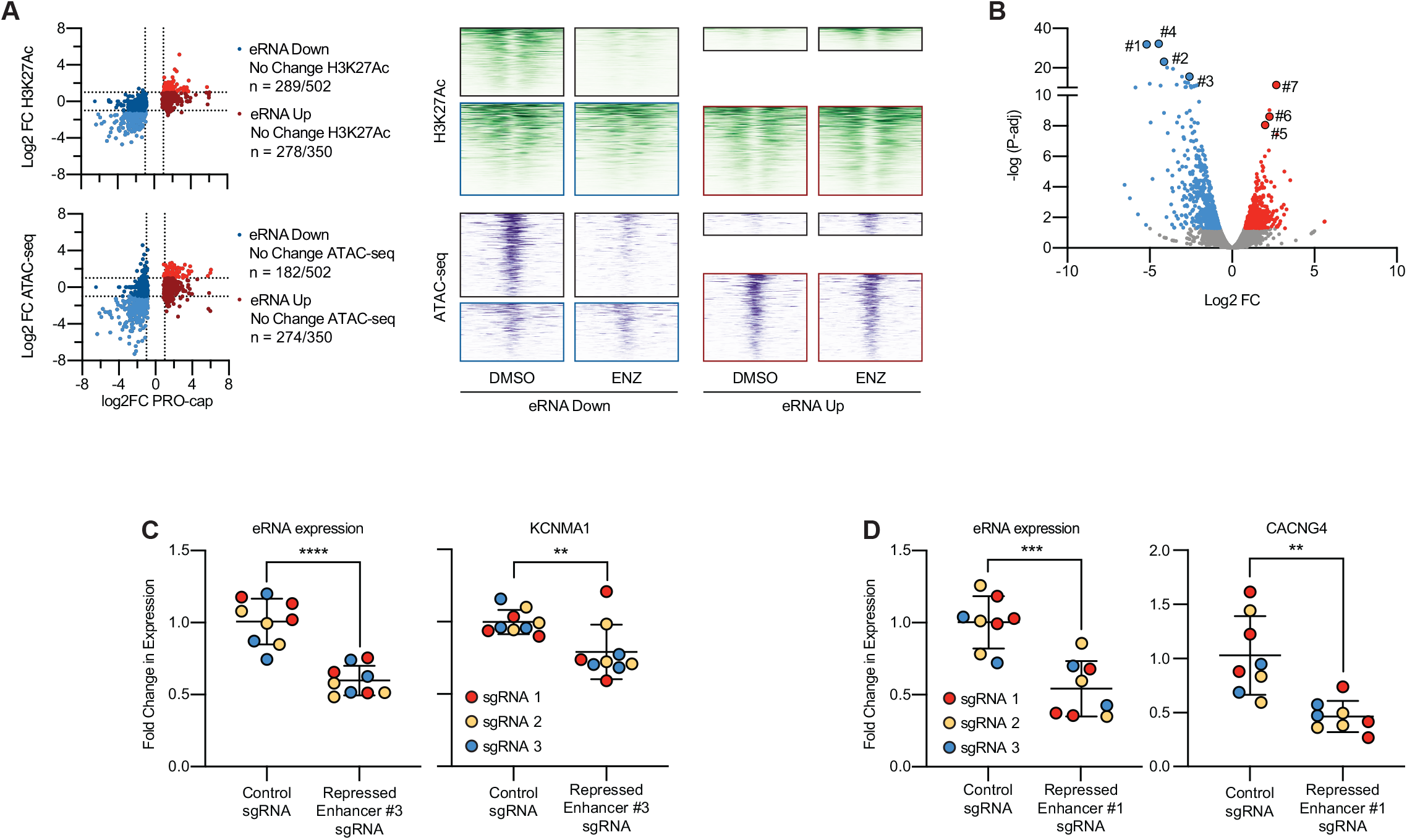
Functional validation of PRO-cap enhancers. **A)** Comparing changes in eRNA expression with ENZ treatment as detected by PRO-cap with changes in H3K27Ac ChIP-seq and ATAC-seq. XY charts and heatmaps show read density 1kb up- and downstream of the center of the ENZ regulated enhancers. Data is from publicly available data(Hwang et al., 2019) (GSE137775). **B)** Selected candidate enhancers for functional analysis. **C)** CRISPRi targeting PRO-cap candidate enhancer #3 with three different sgRNAs significantly reduces expression of the eRNA itself (*P* < 0.0001), and of the downstream target gene *KCNMA1* (*P* = 0.0083). **D)** CRISPRi targeting PRO-cap candidate enhancer #1 with three different sgRNAs significantly reduces expression of the eRNA itself (*P* = 0.0002), and of the downstream target gene *CACNG4* (*P* = 0.0011).

**Figure 4.**
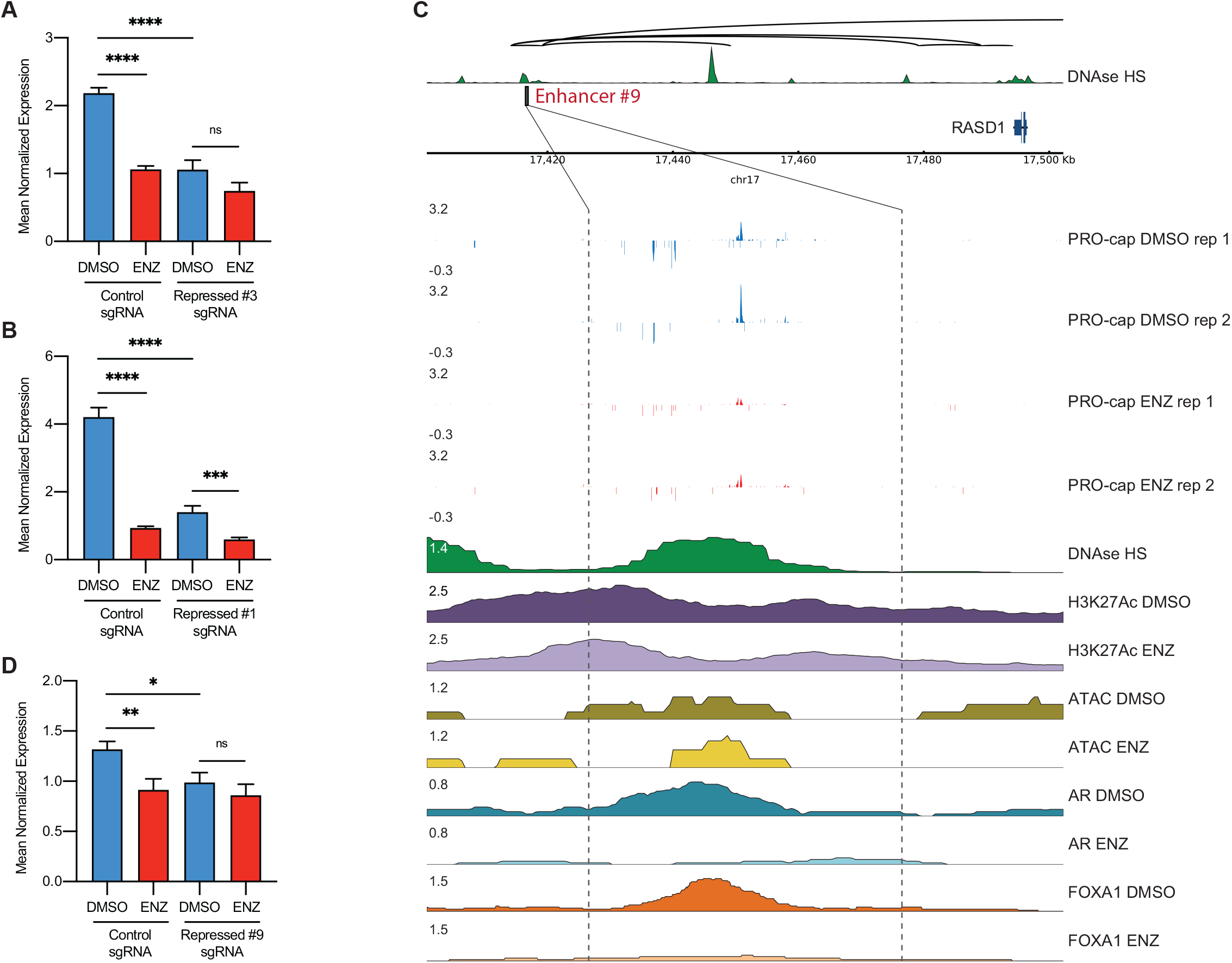
CRISPRi interrogation of PRO-cap enhancer candidates. **A)** CRISPRi targeting PRO-cap candidate enhancer #3 with three different sgRNAs impairs the ENZ-regulation of the downstream target gene *KCNMA1* (2.05-fold change, *P* < 0.0001 vs. 1.42-fold change). **B)** CRISPRi targeting PRO-cap candidate enhancer #1 with three different sgRNAs impairs the ENZ-regulation of the downstream target gene *CACNG4* (4.48-fold change, *P* < 0.0001 vs. 2.35-fold change, *P* = 0.0007). **C)** Shown is the candidate enhancer PRO-cap signal along with published H3K27Ac HiChIP loops(Giambartolomei et al., 2021) and read tracks for DNase (GSM816637), ATAC-seq, and ChIP-seq with and without ENZ for H3K27Ac, AR, and FOXA1 (GSE137775)(Hwang et al., 2019). **D)** CRISPRi targeting PRO-cap candidate enhancer #9 with two different sgRNAs impairs the ENZ-regulation of the downstream target gene *RASD1* (1.44-fold change, *P* = 0.0084 vs. 1.15-fold change).

We next sought to determine whether targeting the TSS altered the ability of the candidate enhancer to regulate gene expression in response to ENZ treatment. To do so we again transfected dCas9-KRAB-expressing LNCaP cells with the chosen sgRNAs followed by 24-hour treatment with ENZ. Again, for both genes we demonstrated significant downregulation with ENZ (*KCNMA1, P* < 0.0001; *CACNG4, P* < 0.0001), with the sgRNA (*KCNMA1, P* < 0.0001; *CACNG4, P* < 0.0001), and a significant reduction in the ENZ downregulation in combination with the sgRNA (*KCNMA1, P* = n.s., 1.42 vs 2.05-fold change; *CACNG4, P* = 0.0007, 2.35 vs 4.48-fold change) (Figure 4A-B).

Next, we functionally validated the ability of PRO-cap to identify changes in enhancer activity that are not detected by ATAC-seq or H3K27ac ChIP-seq assays (Figure 3A). Given that the two above candidates both show concomitant changes in both of those biochemical marks (Figure S3-4), we pursued enhancer candidates which: (1) show significant changes in PRO-cap upon ENZ treatment, (2) show no changes in ATAC-seq or H3K27ac ChIP-seq after ENZ treatment, and (3) are predicted to regulate known ENZ-responsive genes.

The first enhancer candidate (#8) selected resides within an intronic region of *DENND1B* (Figure S5A), a gene that has been shown to be upregulated with ENZ (Figure S5B). While *DENND1B* was not significantly upregulated upon ENZ treatment in control sgRNA-expressing cells, a dramatic increase in its expression was seen in cells expressing sgRNAs against the candidate target #8 with ENZ treatment (*P* < 0.0001, 2.06 vs 0.946-fold change) (Figure S5C). The next enhancer candidate selected (#9) is ∼80 kb upstream of *RASD1* (Figure 4C, Figure S6A), which is a known ENZ-repressed gene (Figure S6B). *RASD1* was significantly downregulated with ENZ treatment in control sgRNA-expressing cells (*P* = 0.0084). Likewise, a significant decrease in RASD1 expression was observed in cells transfected with the candidate #9 targeting sgRNAs than in control sgRNA-expressing cells (*P* = 0.0214). and there was a significant reduction of ENZ downregulation in combination with the sgRNA (*P* = n.s., 1.15 vs 1.44-fold change) (Figure 5D).

Altogether, these *in vivo* functional results confirm the utility of our PRO-cap assay as a highly-sensitive approach to identify putative enhancers and detect changes in their activities genome-wide.

## Discussion

This study highlights the generalizable utility of using eRNA transcription patterns obtained from PRO-cap to detect precise changes in enhancer activity, which has broad applications in human genetics spanning development and disease. Despite previous efforts in generating chromatin accessibility and histone modification landscapes to indirectly map transcriptional regulatory networks, our knowledge of the key regulatory mechanisms that orchestrate the complexity of cellular differentiation, human development, and disease pathogenesis is still limited by our incomplete understanding and characterization of the non-coding genome, in particular with the annotation of cell-state specific transcriptional regulatory elements. Thus, the ability to delineate more precise maps of enhancer activity dynamics will facilitate the systematic examination of the transcriptional programs of developing cells across cell transition states and different differentiation lineages, including disease progression. With a significant fraction of disease-associated risk variants harbored within non-coding regions of the genome, detailed characterization of these regulatory networks that coordinate cell- and tissue-type specificity could provide insights into the molecular mechanisms that underlie dysregulation in numerous disorders and assist in the mapping of variants functional only at specific cellular states. Similarly, precise mapping of enhancer dynamics genome-wide in clinical specimens by PRO-cap can also help better understand disease mechanisms and heterogeneity across patients as well as response and resistance to treatment.

Hence, our study underscores the clinical value of identifying and delineating aberrant distal regulatory elements in cancer to identify potential therapeutic vulnerabilities.

## Materials and Methods

### Cell lines

LNCaP cells (male, ATCC, RRID: CVCL_1379) were maintained in RPMI medium (Gibco, A1049101), supplemented with 10% FBS (Gibco, 10270106), and 1% penicillin-streptomycin (Gibco, 11548876) on poly-L-lysine coated plates. All cell lines were grown at 37 °C with 5% CO_2_. All cell lines were authenticated by STR analysis and regularly tested for mycoplasma.

### ChIP-seq data analysis

For Figure 1B, previously aligned and normalized bigwig files were downloaded from the Cistrome Data Browser allowing for consistent and standardized analysis of data from multiple studies. For other plots previously aligned and normalized bigwig files were downloaded and analyzed in their published format from GEO. Heatmaps and summary plots were generated using the deepTools suite(Ramirez et al., 2016).

### PRO-cap

LNCaP cells were treated with 10 µM enzalutamide or DMSO for 4 or 24 hrs. For PRO-cap, approximately 10 to 30 million cells were processed per sample. Library preparations for two biological replicates each consisting of two technical replicates per condition were processed separately. Cells were permeabilized and run-on reactions were carried out as previously described(Mahat et al., 2016). Following RNA isolation, two adaptor ligations using T4 RNA Ligase 1 (catalog no. M0204; NEB) and reverse transcription using SuperScript III Reverse Transcriptase (catalog no. 18080044; Invitrogen) were performed, with custom adaptors detailed in Table S1. Between adaptor ligations, cap state selection reactions were performed by treating the samples with CIP (catalog no. M0290; NEB) to reduce uncapped RNAs to 5′ hydroxyls and make them incapable of ligating to 5′ adaptor and Cap-Clip (catalog no. C-CC15011H; Cambio) to remove the 5′ cap of transcripts that had undergone guanylation and allow them to be incorporated into the library through 5′ adapter ligation. RNA washes, phenol:chloroform extractions and ethanol precipitations were carried out between reactions. All steps were performed under RNase-free conditions and following manufacturer protocols. Libraries were sequenced on an Illumina HiSeq 3000 following PCR amplification and library clean-up.

### ChIA-PET and Hi-C data processing for downstream gene predictions

Similar to a previously described method(Wang et al., 2018), LNCaP RNA Pol II ChIA-PET interactions(Ramanand et al., 2020) (GSM3423998) were lifted over from GRCh37 to GRCh38. LNCaP Hi-C data (ENCFF676WJO) was downloaded from ENCODE and interactions were lifted over from GRCh37 to GRCh38. ChIA-PET and Hi-C interactions were merged and filtered to remove duplicate interactions. The merged dataset was used as the final 3D interaction set.

Active enhancers and active promoters were inferred based on a merged set of unidirectional and bidirectional PRO-cap peaks from LNCaP treated with either ENZ or DMSO for 4 or 24 hrs. PRO-cap peak regions were intersected with GENCODE (V28) annotated promoters to get a list of active promoter regions. Active enhancers and promoters were overlapped with the final 3D interaction set to identify potential enhancer-gene pairs. A maximum of 1 Mb distance was allowed.

### Analysis of whole genome patient data

Prostate adenocarcinoma whole genome variants were downloaded from TCGA. This included SNV data from 286 samples and indel data from 200 samples after filtering and overlapping with enhancer regions. The effects of these variants were predicted using Funseq2(Fu et al., 2014) (V2.1.4). The motif break and gain events of the variants were extracted from Funseq2 annotations.

### Analysis of PRO-cap data

Differential transcription at eRNA peaks was quantified using EdgeR(Robinson et al., 2010) analysis of the total read counts in the core promoter (-35 to 60 bp from the peak TSS)(Tippens et al., 2020) of the plus and minus strands separately. Induced enhancers were defined as FDR adj p-value < 0.05 in at least one peak direction.

### CRISPRi

Lenti-dCas9-KRAB-blast was a gift from Gary Hon (Addgene plasmid # 89567). Lentivirus was produced in HEK293T cells, and subsequent virus-containing media was used to transduce LNCaP cells, followed by blasticidin selection.

gRNAs against the candidate enhancer regions were designed using CRISPick. (https://portals.broadinstitute.org/gppx/crispick/public)(Doench et al., 2016; Sanson et al., 2018) (Table S1). Custom crRNAs were ordered from IDT and were annealed with Alt-R® CRISPR-Cas9 tracrRNA (IDT, 1072532) to generate sgRNAs according to the manufacturer’s instructions. For each experiment LNCaP-dCas9-KRAB cells were transfected with 30 nM sgRNA complex using Lipofectamine RNAiMAX (Invitrogen, 13778030). 48 hours post-transfection cells were treated with media containing 10 µM enzalutamide or DMSO for a further 24 hours.

### qPCR

RNA was extracted directly from cells using the RNeasy Mini Kit (Qiagen, 74106), and genomic DNA was removed using the DNA-free kit (Ambion, AM1906). RNA was reverse transcribed using random primers and SuperScript IV Reverse Transcriptase (Invitrogen, 18090010). Quantitative real-time PCR was performed on the ViiA 7 system (Applied Biosystems) using HOT FIREPol EvaGreen qPCR mix (Solis Biodyne, 08-24-00020) following the manufacturer’s instruction. Primer sequences are listed in Table S1. All quantitative real-time PCR assays were carried out using three technical replicates using HMBS as the housekeeping gene.

## Supporting information

DataS1-7

## Data availability

PRO-cap enhancer calls are available in Data S1-7. Raw PRO-cap data and bigWig files are available through GEO under accession no. GSE198268. Other datasets used in this study are publicly available under the following accession nos. LNCaP ATAC-seq (accession no. GSE105116). LNCaP DNase-seq (accession no. GSM816637). LNCaP H3K27ac ChIP-seq (accession no. GSE107780). LNCaP H3K4me1 ChIP-seq (accession no. GSE73783). LNCaP RNA Pol II ChIA-PET (accession no. GSM3423998). LNCaP Hi-C (accession no. ENCFF676WJO). LNCaP AR ChIP-seq, LNCaP FOXA1 ChIP-seq, and LNCaP H3K27ac ChIP-seq (accession no. GSE85558). ATAC-seq, AR ChIP-seq, FOXA1 ChIP-seq, and H3K27ac ChIP-seq (accession no. GSE137775) of LNCaP treated with ENZ. ASH2L ChIP-seq (accession no. GSE60841) of VCaP treated with R1881 or cultured in media containing CS FCS. ChIP-seq (accession nos. GSE137775, GSE125245, GSE94682, GSE70079, GSE40269, GSE28264, and GSE80256) of LNCaP treated with ENZ, R1881, DHT, or cultured in media containing CS FCS. ChIP-seq (accession no. GSE130408) of normal prostate tissue, primary prostate cancer, and metastatic prostate cancer. LNCaP STARR-seq (GSE151064) of AR binding regions treated with DHT or EtOH.

## Code availability

PRO-cap enhancers were called using Peak Identifier for Nascent Transcript Starts (PINTS)(Yao et al., 2022). All analysis was performed using common publicly available tools.

## Figures

**Figure S1.**
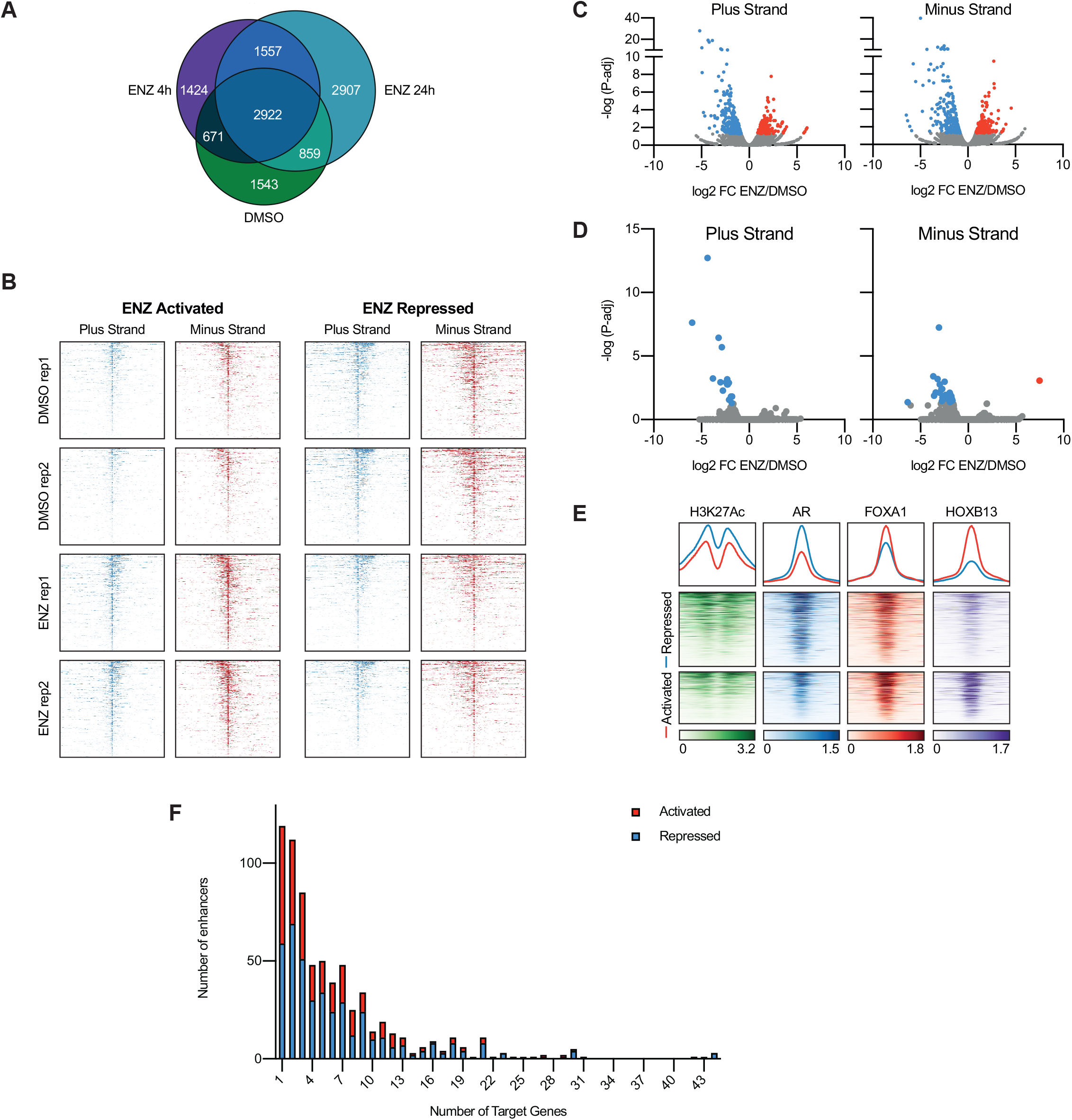
PRO-cap sequencing of eRNAs identifies enhancers affected by enzalutamide. **A)** Venn diagram of the overlap of enhancer regions between the three treatments paradigms **B)** Heatmap of the PRO-cap signal 250 bp up- and downstream of ENZ activated or repressed peaks from two replicates of LNCaP cells treated with DMSO or 10 µM ENZ for 24 hours. **C)** PRO-cap analysis of LNCaP cells treated with 10 µM enzalutamide (ENZ) for 24 hours identified 853 putative enhancers which demonstrated significant differential activation or repression on the plus or minus strands as determined by edgeR analysis (FDR < 0.05). **D)** PRO-cap analysis of LNCaP cells treated with 10 µM enzalutamide (ENZ) for 4 hours identified 44 putative enhancers which demonstrated significant differential activation or repression on the plus or minus strands as determined by edgeR analysis (FDR < 0.05). **E)** Heatmaps and summary plots show read density 1kb up- and downstream of the center of the ENZ regulated enhancers in metastatic PCa. Data is from publicly available ChIP-seq data(Pomerantz et al., 2020) (GSE130408) and is displayed as the mean signal from n = 8 metastatic PCa samples. **F)** Histogram demonstrating the number of downstream target gene predictions for the 853 ENZ-regulated enhancers identified with PRO-cap.

**Figure S2.**
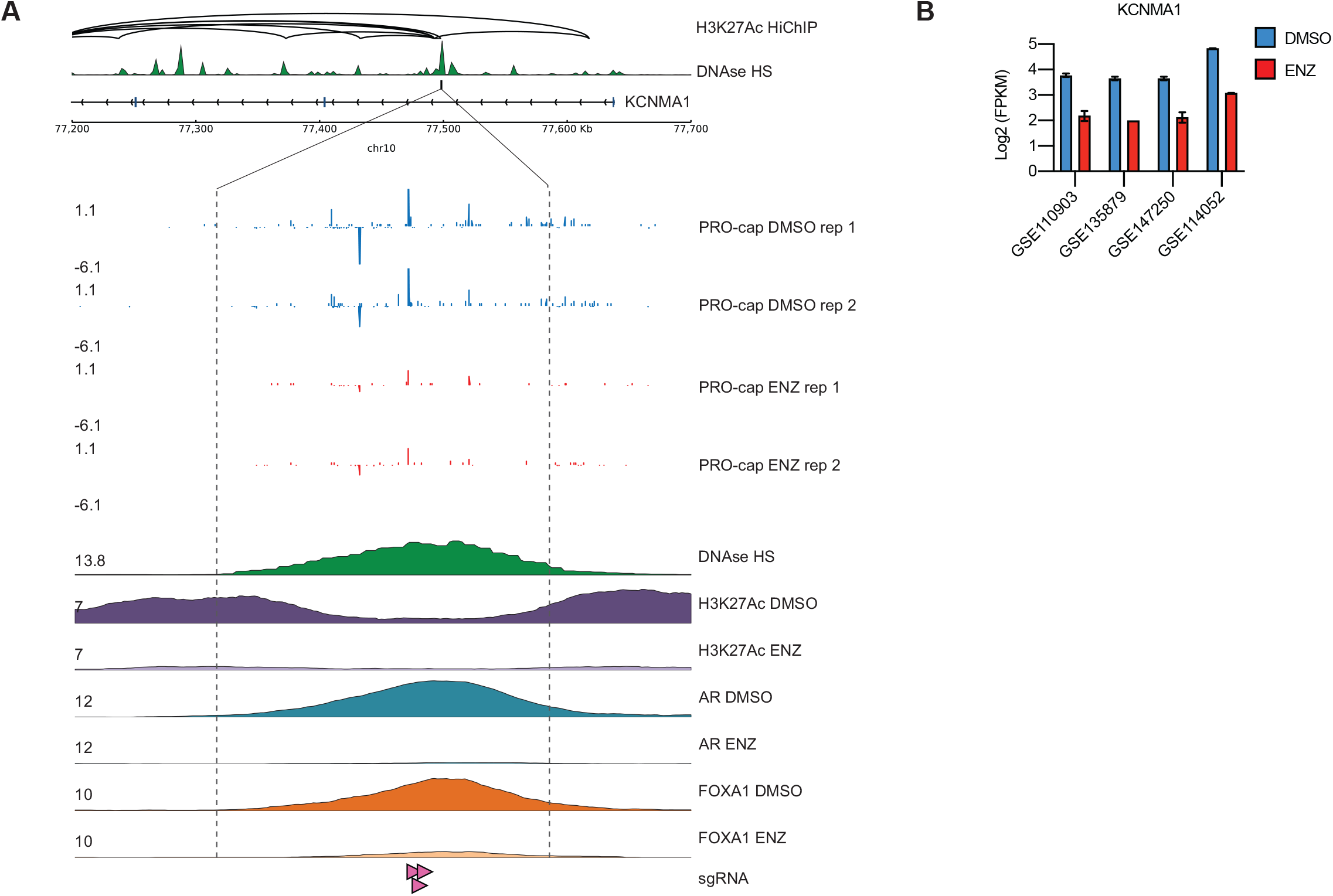
PRO-cap helps prioritize candidate enhancer regions to search for cancer-associated germline and somatic variants. **A)** Analysis of 286 PCa whole genomes identified 137 single nucleotide variants (SNV) in the ENZ-regulated enhancers identified via PRO-cap, 20 of which were recurrent in more than one patient. **B)** Example of an enhancer downregulated by ENZ which demonstrates two separate recurrent SNVs, one in two patients that breaks an ESRRA motif, and another in two different patients that creates an HNF4 motif. **C)** Example of a PCa risk variant(Conti et al., 2021) which overlaps with an enhancer identified via PRO-cap

**Figure S3.**
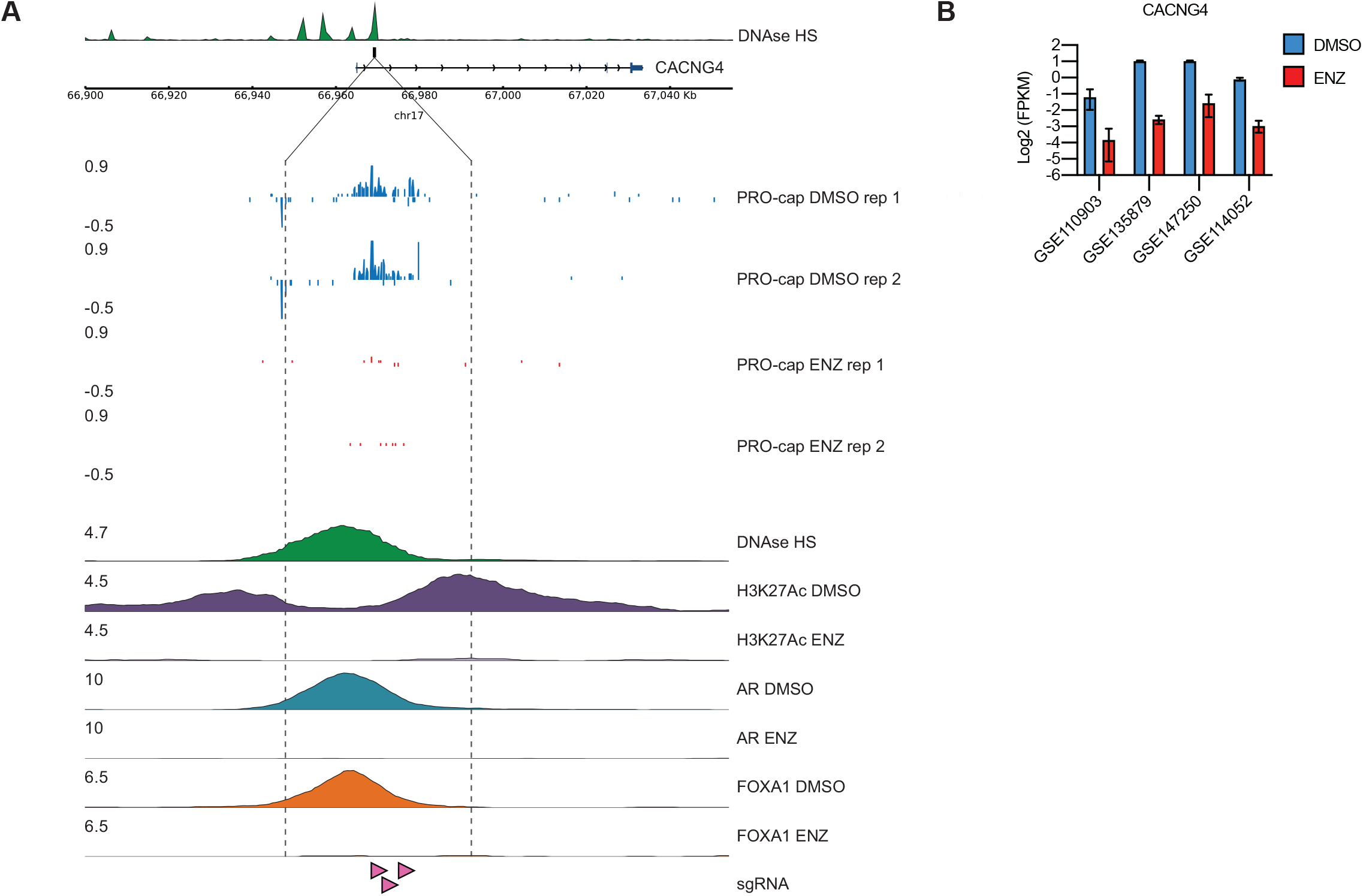
Candidate Enhancer #3. **A)** Shown is the candidate enhancer PRO-cap signal along with published H3K27Ac HiChIP loops(Giambartolomei et al., 2021) and read tracks for DNase (GSM816637), and ChIP-seq with and without ENZ for H3K27Ac, AR, and FOXA1 (GSE137775)(Hwang et al., 2019). **B)** Expression of the candidate downstream target gene *KCNMA1* in response to ENZ from multiple published RNA-seq data sets, accession number as indicated.

**Figure S4.**
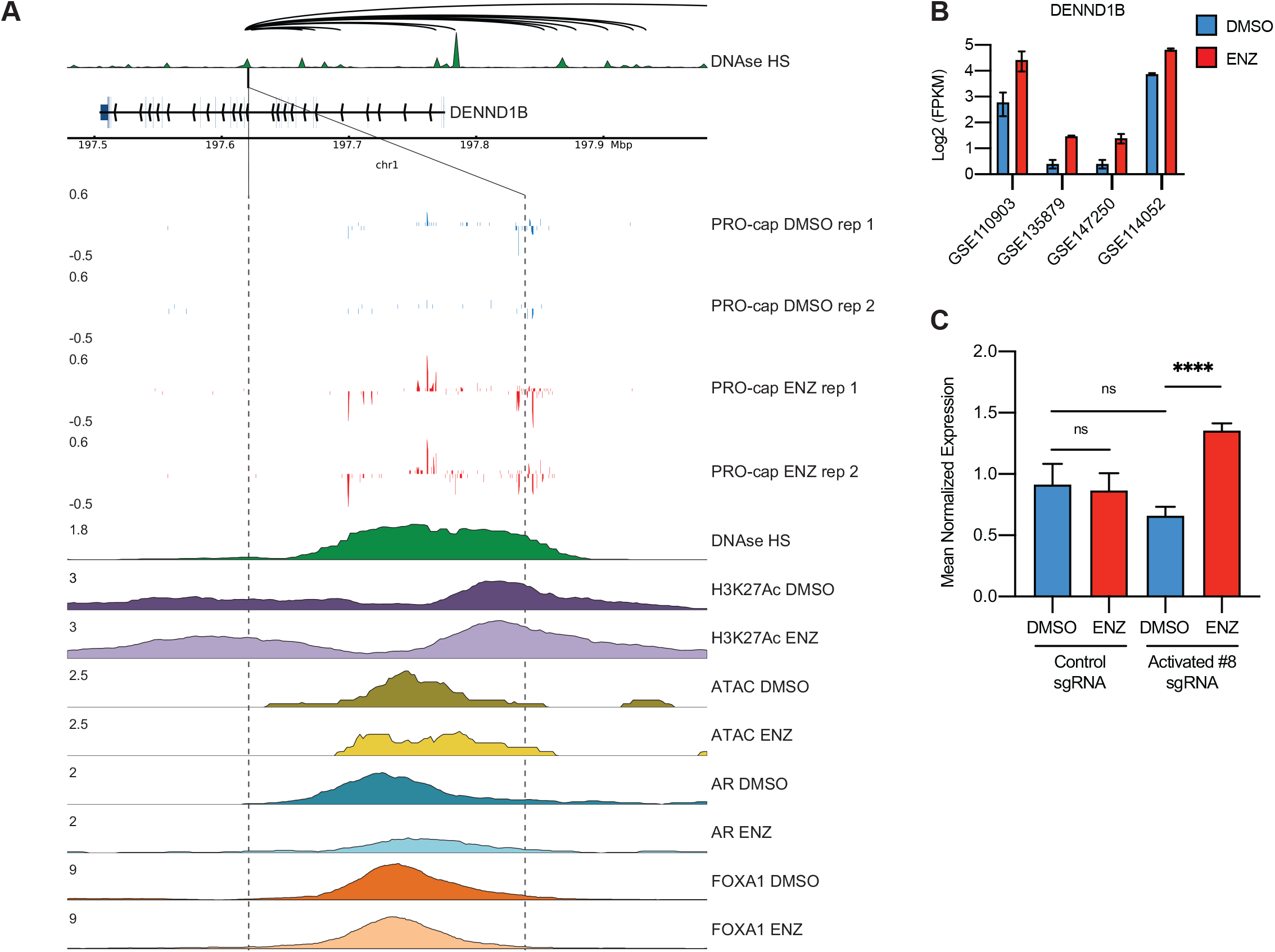
Candidate Enhancer #1. **A)** Shown is the candidate enhancer PRO-cap signal along with published H3K27Ac HiChIP loops(Giambartolomei et al., 2021) and read tracks for DNase (GSM816637), and ChIP-seq with and without ENZ for H3K27Ac, AR, and FOXA1 (GSE137775)(Hwang et al., 2019). **B)** Expression of the candidate downstream target gene *CACNG4* in response to ENZ from multiple published RNA-seq data sets, accession number as indicated.

**Figure S5.**
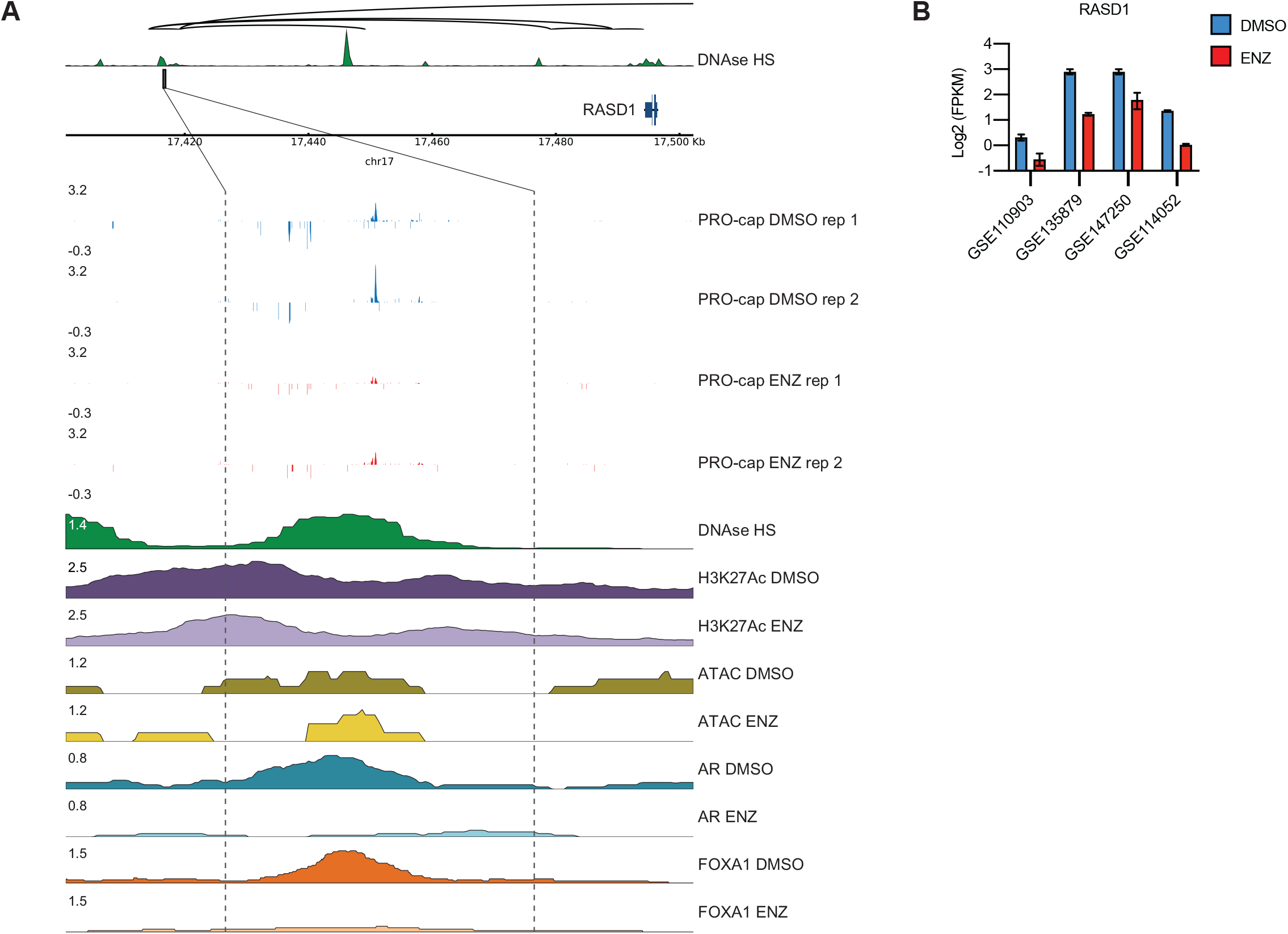
Candidate Enhancer #8. **A)** Shown is the candidate enhancer PRO-cap signal along with published H3K27Ac HiChIP loops(Giambartolomei et al., 2021) and read tracks for DNase (GSM816637), ATAC-seq, and ChIP-seq with and without ENZ for H3K27Ac, AR, and FOXA1 (GSE137775)(Hwang et al., 2019). **B)** Expression of the candidate downstream target gene *DENND1B* in response to ENZ from multiple published RNA-seq data sets, accession number as indicated. **C)** CRISPRi targeting PRO-cap candidate enhancer #8 with three different sgRNAs impairs the ENZ-regulation of the downstream target gene DENND1B (0.946-fold change vs. 2.06-fold change P < 0.0001).

**Figure S6.**
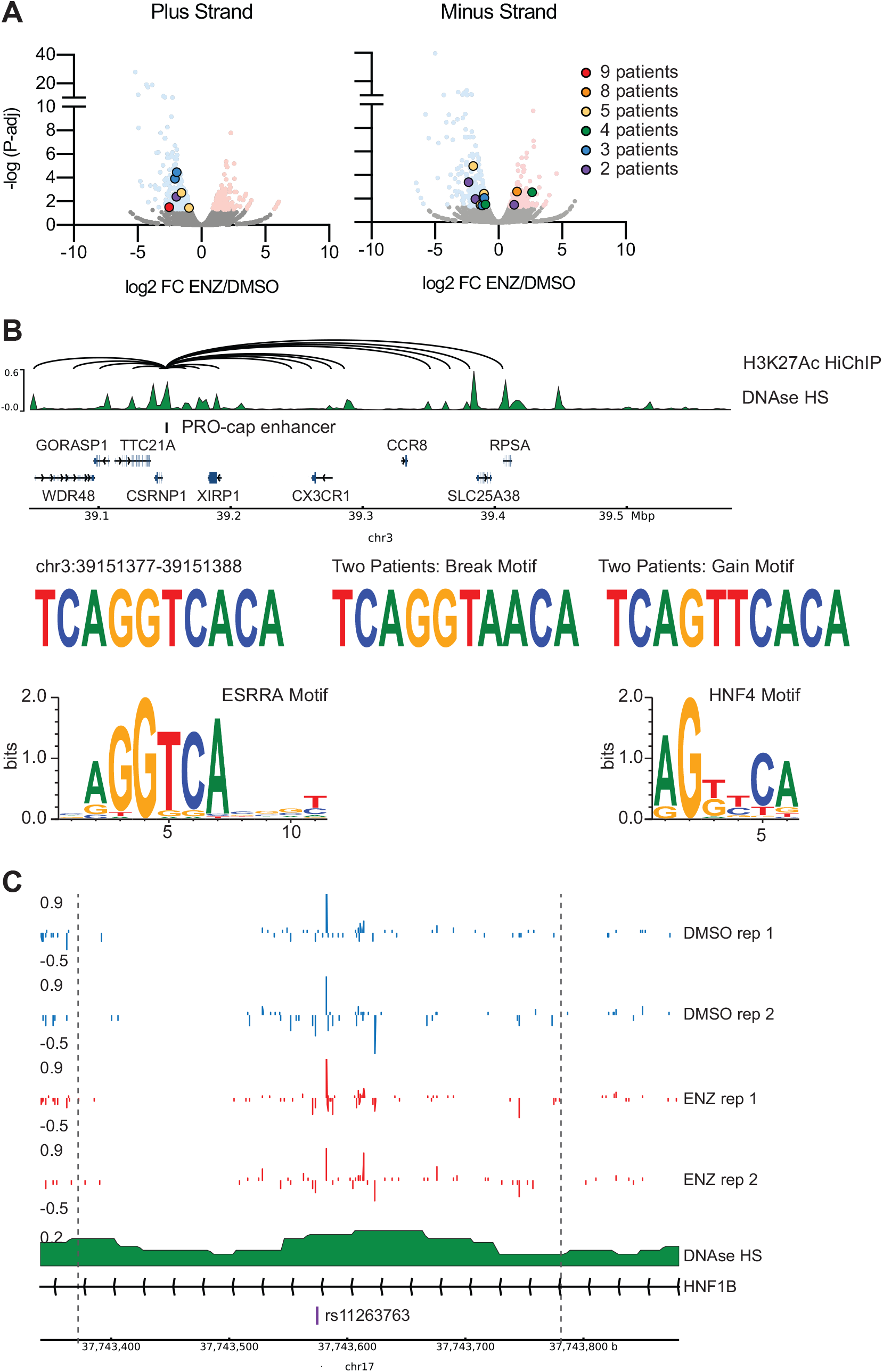
Candidate Enhancer #9. **A)** Shown is the candidate enhancer PRO-cap signal along with published H3K27Ac HiChIP loops(Giambartolomei et al., 2021) and read tracks for DNase (GSM816637), ATAC-seq, and ChIP-seq with and without ENZ for H3K27Ac, AR, and FOXA1 (GSE137775)(Hwang et al., 2019). **B)** Expression of the candidate downstream target gene *RASD1* in response to ENZ from multiple published RNA-seq data sets, accession number as indicated.

